# Spatially explicit modelling of kelp-grazer interactions: a seascape approach for effective habitat enhancement using artificial reefs

**DOI:** 10.1101/2020.06.23.166090

**Authors:** Ferrario Filippo, Thew Suskiewicz, Ladd Erik Johnson

## Abstract

1. Artificial structures are sprawling along the coast affecting the aspect and the functioning of shallow coastal seascapes. For years, the ecology of artificial structures has been investigated mainly in contrast to natural coastal habitats. However, it is increasingly emerging that structuring processes, such as trophic interactions, can depend on properties of the surrounding landscape. Heterogeneity of coastal seafloor and habitats could thus play a major role in determining the variability of ecological outcomes on artificial structures. Artificial reefs are being used in coastal areas in attempts to restore and enhance marine habitats and communities, including large brown seaweed (“kelp”), to offset habitat loss and mitigate coastal development impacts. The outcome of enhancement projects using artificial reefs have not always been either consistent or positive. Overlooking the effect of strong ecological interactions adds a high level of uncertainty and can undermine the success of these efforts. In Eastern Canada, top-down control exerted by green sea urchins (*Strongylocentrotus droebachiensis*) can seriously compromise the success of artificial reefs for kelp enhancement. Importantly, urchin interactions with macroalgae are likely to be influenced by the bottom composition. A seascape approach could thus integrate behavior and habitat heterogeneity.
2. We investigated whether the local seascape could create zones of differential grazing risk for kelp outplanting kelp (*Alaria esculenta*) on artificial blocks on an heterogenous bottom. Adopting a spatially explicit framework, we determined how seascape affected the urchin use of the habitat and used this information to map the grazing risk throughout the area.
3. Kelp survival was a function of frequency of urchin presence throughout the study site. While urchins avoided sandy patches, bottom composition and algal cover modulated the within-patch urchin use of the habitat. This translated in the heterogeneity of grazing risk intensity.
4. *Synthesis and applications*. The presence of discrete seascape features locally increased the grazing risk for kelp by differentially affecting the urchin’s usage of the habitat, even within the same bottom patch. Incorporating this information when planning artificial reefs could minimize the detrimental grazing risk thus increasing the rate of success and ensuring lasting results.

## Introduction

The ecological functioning of coastal artificial structures and their quality as a novel habitat for marine species have been an active research topic for almost three decades now, and technological advances, general frameworks and syntheses are being proposed to integrate ecological concepts in coastal development and foster its sustainability (Bishop et al., 2017; Dyson & Yocom, 2015; Mayer-Pinto et al., 2017; Perkol-Finkel, Hadary, Rella, Shirazi, & Sella, 2018). However, whether investigating biodiversity, colonization and ecological succession, community ecology, or species genetic diversity, so far the predominant approach to study artificial structures has been framed in the dualism typically contrasting alternative categories, such as artificial-natural, vegetated-unvegetated, hard-soft bottoms, impacted-pristine habitats. While these binary contrasts account for a relevant part of the observed differences between systems, the presence and the intensity of structuring ecological processes can critically determine the variability of the biological communities on artificial structures. For example, recognizing the effect of trophic interactions and biotic disturbance on the survival of macroalgae can help understanding the composition of the algal community on coastal defence structures (Ferrario, Iveša, Jaklin, Perkol-Finkel, & Airoldi, 2016; Perkol-Finkel, Ferrario, Nicotera, & Airoldi, 2012).

Interspecific ecological interactions are affected by the characteristics of the surrounding landscape which influence how species use their habitat. These landscape ecology concepts are well exemplified in the terrestrial literature. For instance, caribous in boreal regions are more likely to roam and graze in their preferred resource patches the closer these are to refuge-offering forest stands (Mason & Fortin, 2017), similarly the wolf use of the landscape modulated the relative preference of elks for different land-cover types (Fortin et al., 2005). In western North America, the spread of aggressive mountain pine beetles and eruptions of infestation spots is influenced by landscape properties, namely tree species composition and host density, that determine heterogeneous forest patches offering differential movement resistance and high mobility corridors (Powell, Garlick, Bentz, & Friedenberg, 2018).

Similarly, the role of seascape – a spatially heterogeneous area of coastal environment perceived as a mosaic of patches (Boström, Pittman, Simenstad, & Kneib, 2011) – in affecting structuring ecological processes has been increasingly emerging. For example, Foster (1990) highlighted how failing to recognize coastal sites’ heterogeneity challenged the ecologist ability to generalize mechanisms underlying species’ zonation over wider geographical scales than those at which observations were made. Still resorting to contrasts in factorial designs, Micheli & Petterson (1999) showed the role of vegetated bottom patches as corridors for predators, so that oyster reefs at different distances from salt marsh edges were equally predated by blue crabs only in the presence of a seagrass matrix. Despite an overall lack of seascape maps of proper resolution and the inability to track animals compared to their terrestrial counterpart, marine ecologists are increasingly considering the spatially explicit seascape configuration to better resolve the mechanisms underlying the ecological processes. For example, predation risk for Mediterranean sea urchin *Paracentrotus lividus* and predator identity emerged as a function of the arrangement of seagrass patches (Farina et al., 2016). Aerial imagery in clear tropical water systems allowed to show that grazing pressure on macroalgae decreases with the distance from the nearest coral reef providing shelter to fish (Gil, Zill, & Ponciano, 2017; Madin, Madin, & Booth, 2011).

Understanding the effects of seascape in modulating the intensity of ecological structuring processes is a pivotal element to better forecast the ecological trajectory of marine artificial structures, considering the local deployment setting to go beyond the contrast of natural and artificial habitats. Such an ability will greatly assist coastal managers and conservationists in minimizing the risk of failure, increasing the consistency of outcomes and ultimately the sustainability of those habitat enhancement projects resorting on artificial structures. To this regard, the eastern Canadian coastal ecosystems and in particular the Gulf of Saint Lawrence (GSL) represent a case study for the use of artificial reefs.

The Canadian environmental policy requires remediation efforts to offset damages to marine habitats caused by coastal development, with a particular focus on protecting or creating new fish habitat (Fisheries Act, 2019). Fostered by this legislation, several artificial reefs have been employed by different project proponents to offset the loss of habitat. Artificial reefs are a well-studied approach for habitat restoration in both tropical and temperate ecosystem (Fabi et al., 2011; Feary, Burt, & Bartholomew, 2011) and modular design – i.e. reefs composed of small independent units like concrete blocks – is becoming more frequent (Dyson & Yocom, 2015; Tessier et al., 2015). In Canada, modular reefs have been built in the attempt to provide a suitable substratum for large brown seaweed canopies, commonly known as kelp, and to create new kelp populations (Department of Fisheries Oceans, personal communication).

The deployment of artificial reefs targeting kelp, do not merely represent a compliance with the law but a real opportunity to actively enhance kelp presence and distribution in the GSL region. Indeed, the high ecological value of kelp makes them one of the most desirable habitats in coastal environments (Steneck et al., 2002), yet the GSL is characterized by a set of environmental and biological conditions which confine kelp habitats into a very narrow and shallow (< 3 meters) subtidal refugia. Indeed, while ice scouring during the winter can seriously damage kelp in the intertidal and shallow subtidal (P. Gagnon, Himmelman, & Johnson, 2004; Keats, 1991), the main threat to kelp beds come from overgrazing by the green sea urchin *Strongylocentrotus droebachiensis* (O.F. Müller, 1776). This echinoderm can locally reach densities of 100s individuals per meter square (P. Gagnon et al., 2004) and maintain the system locked in a “barren” state dominated by crustose coralline algae (CCA) (Ling et al., 2015; Steneck, Leland, McNaught, & Vavrinec, 2013), far less diverse and productive than kelp dominated systems.

Overgrazing by sea urchins is thus one cause of failure of major concern for kelp enhancement via artificial reefs in the GSL. However, as sea urchins are mainly associated with rocky bottoms, the heterogeneity of the seascape (i.e. the extent and the spatial arrangement of patches of sand, gravel and cobbles, boulders, and bedrock) could potentially offer kelp with refugia from herbivory.

In this work we 1) investigated whether examination of seascape heterogeneity could identify areas of lower risk (i.e. refugia) for kelp transplanted onto artificial reef modules along a heterogeneous bottom, and we 2) characterized which seascape properties define these refugia. In particular, we focused our attention on the effect of three main groups of seascape properties: bottom composition, biological factors (e.g. algal presence), and spatial arrangement (e.g. distances). Several *a priori* predictions can be made. We anticipate that the grazing pressure by *S. droebachiensis* on kelp will be high in areas with high hard bottom percent cover, while grazing risk will decrease as soft bottom cover increase. Biological factors, such as the presence of other algae, could either attract urchin and thus increase the exposure of kelp to grazing, or alternatively, divert urchin feeding activity thus increasing kelp survival. Finally, we predicted that urchin movement could potentially be affected by the spatial configuration of the artificial reef modules, thus we also wanted to understand whether artificial blocks would favor urchin movement (i.e. stepping stones) or act like local sinks and retain urchins.

## Methods

### Study Area

The study was conducted at the Baie-du-petit-Métis (48°40’37.30” N, 68° 0’41.24” W; Québec, Canada) along the south shore of the St Lawrence Estuary (Fig. 1). On the northwest side, the bay is separated from the St Lawrence River by a narrow peninsula characterized by rocky shores, while sandy beaches become predominant moving counterclockwise towards the eastern shore. A wide intertidal mudflat with sparse rocky outcrops is followed subtidally by areas of soft sediments, gravel and boulders heterogeneously arranged. Fucoid seaweeds (i.e. *Ascophyllum* and *Fucus* spp.) are the most abundant vegetation on the rocky intertidal. Kelp presence in the bay is spatially confined to rocky intertidal pools and the very shallow subtidal (∼1m) and is composed of just a few species: *Alaria esculenta, Saccharina latissima*, and *Agarum clathratum*.

**Figure 1.**
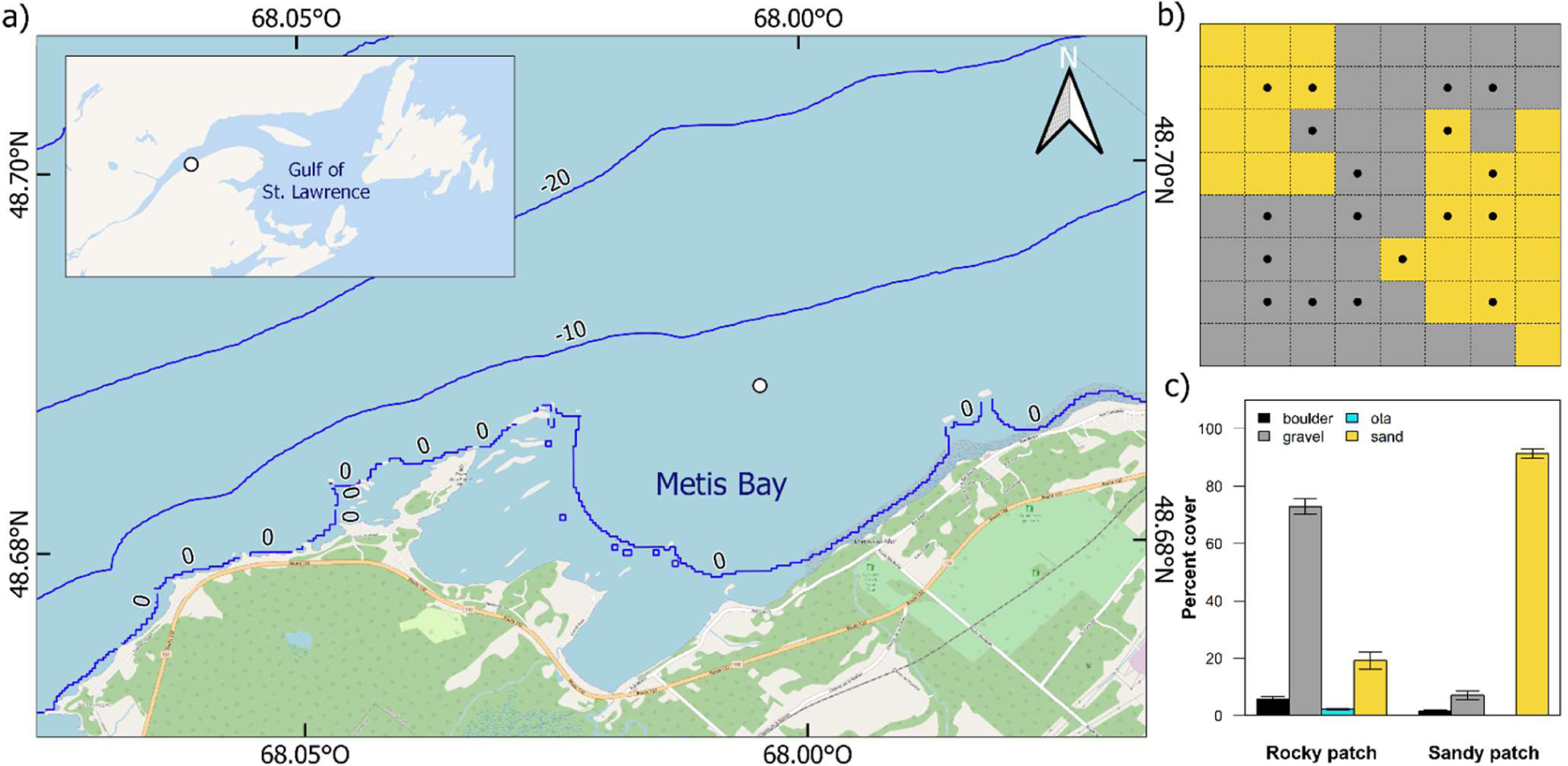
Map of the Metis bay, Spatial configuration and bottom composition and of the study site. a) Location of the study site in the Metis Bay and regional context (open circle). Isobaths represented as blue lines at 10 m depth intervals. Basemap acquired from OpenStreetMap. b) The study site included two sandy patches (yellow cells) and one rocky patch (grey cells). Black dots represent position of artificial blocks with transplant at the beginning of the experiment. Dashed lines depict cells borders. b) Average percent cover of boulders, gravel, sand and algae (ota) of cells within rocky and sandy patch (mean ± SE).

### Experimental design and modelling approach

We wanted to map the grazing risk for kelp within a heterogeneous area. We thus identified a site featuring a heterogeneous bottom (i.e. a mix of sandy and rocky patches) over a continuous squared area of 256 m^-2^ at a depth of 8 m meter within the bay (Fig. 1b). The site was mapped to acquire relevant seascape variables while the survival of kelp outplanted on artificial blocks was monitored throughout the summer. We then adopted a two-step modeling approach to describe first the survival of kelp as a function of a relevant urchin metric (e.g. presence/absence, density), and second the habitat use by sea urchins as a function of seascape properties. By coupling the two models we thus obtained a map of the grazing risk in the area.

### Bottom mapping

We mapped the study site by means of photographic sampling performed by SCUBA divers over a permanent grid of 16×16 m divided in 64 cells of 2×2m size (Fig. S1). To trade-off the acquisition of detailed pictures and the image footprint, each cell was virtually divided in four parcels of 1 m side, each of which was photographed using a GoPro Hero3^+^ camera mounted on a PVC frame with a squared base (0.8×0.8 m, Fig S2). We defined the area extent *a priori* to optimize the workload (*Supporting Information – Bottom mapping*).

Photographs were analyzed to quantify percent cover of substrate type and selected algal categories (*Supporting Information – Bottom mapping*). Substrate type included three categories: “boulders” (BLD, >25cm), “cobbles and gravel” (CAG, 0.2-25 cm) and “sand” (SND, < 0.2 cm). Cobbles and gravel were grouped in the same category to facilitate the analysis. Algal categories included foliose algae (‘Algae’, mainly red algae), turf and sediment (‘TurfSed’), and crustose coralline algae (CCA). Algal cover was estimated at all sampling times to monitor changes, while bottom type classification was conducted only on photographs taken at the beginning of the experiments and then assumed to remain constant.

### Kelp survival experiment and urchin distribution

We outplanted one reproductive thallus of *A. esculenta* on each of 18 artificial blocks (cinder blocks, 15×40×20 cm, Fig. S3) on August 5^th^ 2015 (T_days_=0) and we assessed kelp survival periodically, every 10 to 20 days depending on weather conditions, through the summer until October 13^th^ (T_days_=69). Kelp survival was assessed again almost one year after. We positioned the blocks in the center of randomly chosen grid cells among those that had eight adjacent cells to allow the calculation of neighboring statistics. We sourced kelp from a natural population occurring either in nearby tidal pools or in the shallow subtidal and outplanted the sporophytes according to Carney et al. (2005, Supporting Information).

We considered one kelp as “survived” whenever its meristem was present (i.e. when the length of the thallus was equal to or greater than the length of the stipe between the holdfast and the beginning of the meristem (Mann, 1973)) and as “lost” otherwise or when the kelp was missing.

We counted the number of visible urchins in all photographs taken at each sampling time using the Cell Counter plug-in for ImageJ 1.47, considering only individuals with a test diameter greater than 20 mm. Divers additionally recorded the number of urchins found in the artificial block cavities (hereafter “hiding urchins”). We assessed hiding urchins only for cells containing artificial blocks (i.e. one value per cell).

### Verification of outplant technique and suitable kelp growing conditions

Grazing is not the only possible cause for kelp loss, particularly with transplant experiments. Poor growing conditions, transplant stress and dislodgement are other possible vectors leading to the loss of individual plants. To verify our transplant technique and test the suitability of the physical environment we also transplanted kelp onto a frame suspended off the bottom (i.e. three ropes arranged in a pyramidal structure, Supporting Information) and largely immune from urchin grazing. We deployed the kelp frame in July 2015 and populated the lines with kelp plants spaced 30 cm apart. In October we recovered all kelp transplants, recording for each i) its position (depth) on the frame, ii) whether it had “survived” (described above), iii) whether or not transplants developed reproductive sporophylls and, if so, iv) their length.

### Statistical analyses

#### Kelp survival model

We modeled kelp survival as a function of different sea urchins related metrics using survival analysis, that models the “time until an event occurs” – i.e. survival time - and accommodates censoring retaining individuals with unknown survival time in the dataset (Kleinbaum & Klein, 2005). We defined the survival time as the number of days passed from T_days_=0 to the midpoint of the interval between two consecutive sampling dates within which the event ‘loss of one kelp’ occurred (i.e. survival time = 7 for an event occurred between T_days_=0 and T_days_=14). We considered six predictor variables: ‘urchin frequency’ defined as the frequency of times when urchins were present in a cell with a block (FreqLocal), ‘urchin density’ as the median of the average urchin density recorded at each sampling time in a cell with a block (DensLocal), two predictors describing the median of either the urchin frequency or the medians of average urchin density in the eight cells surrounding to the focal one (FreqNeib and DensNeib respectively), ‘frequency of hiding urchins’ as the frequency of times when urchins were found hiding in one block (HidFreq), and finally the median of the number of urchins hiding in one block (HidNum). Only data from 2015 were analyzed.

We fitted semi-parametric Cox Proportional Hazards (CoxPH), Weibull and exponential parametric survival models for each predictor and a null hypothesis, and compared the 21 resulting candidate models via AICc (Second-order Akaike Information Criterion for small sample sizes). We checked the assumption requirements and reported the results only for CoxPH models since they always outcompeted their parametric alternatives (Table 1, Supporting Information).

**Table 1.** Kelp Survival model selection. Model selected is in bold ΔAIC < 2).

| COX models |  |  |  |  |
| --- | --- | --- | --- | --- |
| Survival ~ ... | AICc | $\Delta AICc^*$ | Akaike weights<sup>§</sup> | rank |
| null (i.e. ~1) | 55.7 | 18.89 | 0.000 | 7 |
| DensLocal | 39.9 | 3.07 | 0.176 | 2 |
| DensNeib | 51.3 | 14.46 | 0.001 | 5 |
| <u>FreqLocal</u> | **36.9** | **0.00** | **0.821** | **1** |
| FreqNeib | 51.0 | 14.17 | 0.001 | 4 |
| <u>HidNum</u> | 50.1 | 13.29 | 0.001 | 3 |
| HidFreq | 53.9 | 17.01 | 0.000 | 6 |

\* difference between AICc of a given model and the AICc of the best model.
<sup>§</sup> ratio of the $\Delta AICc$ of a given model relative to the whole set of candidate models. Higher scores indicate the probability that the model is the best among the whole set of candidate models (Mazerolle, 2006).

#### Urchin use of the habitat

We modeled the sea urchin presence/absence as a function of four major seascape properties: bottom composition, biological factors, spatial configuration and spatial arrangement of the artificial reef (i.e. reef setup). In addition to the variables described in the bottom mapping section, predictors included the median in neighboring parcels of CAG (CAGNeib), Algae (AlgaeNeib), CCA (CCANeib) and of TurfSed (TurfSedNeib). We assigned each parcel to either a sandy or a rocky patch (*Supporting Information – bottom mapping*) and calculated spatial configuration predictors as the shortest distance of a parcel from a sandy patch (dSND), from a rocky patch (dRCK), and from the closest parcel containing a boulder (dBLD). Finally, we considered the reef setup calculating the distance from the closet parcel of a cell containing a block. All distances were in relative unit (i.e. between parcels distance).

We checked all potential predictors for collinearity and retained those deemed more relevant. We modeled urchin presence in each parcel at each sampling time using Binomial Generalized Linear Mixed Models with a logit link and “parcel” as a random effect following Zuur et al. (2009); percent covers predictors were arcsin transformed. We built a set of 17 candidate models that included selected combinations of seascape properties (Table 2) and averaged the those having a ΔAIC ≤ 2 to decrease the selection bias (Mazerolle, 2006). See *Supporting Information* for more details on analyses.

**Table 2.** Urchin use of the habitat model selection and averaging. Models selected for averaging are in bold (ΔAIC < 2). Sub-global models are named “Glb”.

| Predictors |  |  |  |  |  |  |  |  |  |  |  |
| --- | --- | --- | --- | --- | --- | --- | --- | --- | --- | --- | --- |
| Model | Bottom (% cover) | | Biological (% cover) | | | Spatial configuration | | Reef setup | $\Delta AIC^*$ | Akaike weights<sup>§</sup> | rank |
| *Bottom + Spatial + Setup* |  |  |  |  |  |  |  |  |  |  |  |
| Glb1 | BLD | CAG |  |  |  | dSND | dB LD | dBLOCK | 4.56 | 0.033 | 7 |
| M1 | BLD | CAG |  |  |  | dSND | dB LD |  | 3.92 | 0.045 | 5 |
| M2 | BLD | CAG |  |  |  |  |  | dBLOCK | 4.35 | 0.036 | 6 |
| M3 | BLD | CAG |  |  |  |  |  |  | 3.04 | 0.070 | 4 |
| *Bottom + Biological + Setup* |  |  |  |  |  |  |  |  |  |  |  |
| **Glb2** | **BLD** | **CAG** | **Algae** | **AlgaeNeib** | **CCA** |  |  | **dBLOCK** | **0.98** | **0.195** | **3** |
| **M4** | **BLD** | **CAG** | **Algae** | **AlgaeNeib** | **CCA** |  |  |  | **0.10** | **0.303** | **2** |
| M5 |  |  | Algae | AlgaeNeib | CCA |  |  | dBLOCK | 43.00 | 0.000 | 14 |
| M6 |  |  | Algae | AlgaeNeib | CCA |  |  |  | 41.64 | 0.000 | 13 |
| M7 |  |  | Algae |  | CCA |  |  |  | 46.82 | 0.000 | 15 |
| **M8** | **BLD** | **CAG** | **Algae** |  | **CCA** |  |  |  | **0.00** | **0.318** | **1** |
| M9 |  |  | Algae |  | CCA |  |  | dBLOCK | 48.43 | 0.000 | 16 |
| *Biological + Spatial + Setup* |  |  |  |  |  |  |  |  |  |  |  |
| Glb3 |  |  | Algae | AlgaeNeib | CCA | dSND | dB LD | dBLOCK | 16.09 | 0.000 | 10 |
| M10 |  |  | Algae | AlgaeNeib | CCA | dSND | dB LD |  | 16.03 | 0.000 | 9 |
| M11 |  |  |  |  |  | dSND | dB LD | dBLOCK | 33.04 | 0.000 | 12 |
| M12 |  |  |  |  |  | dSND | dB LD |  | 32.51 | 0.000 | 11 |
| M13 |  |  | Algae |  | CCA | dSND | dB LD |  | 15.05 | 0.000 | 8 |
| M14 |  |  |  |  |  |  |  | dBLOCK | 104.79 | 0.000 | 17 |

\* difference between AIC of a given model and the AIC of the best model.
§ ratio of the $\Delta AIC$ of a given model relative to the whole set of candidate models. Higher scores indicate the probability that the model is the best among the whole set of candidate models (Mazerolle, 2006).

All statistical analyses were performed using R 3.3.2 (R Core Team, 2016). We used packages ‘*raster’* for spatial analysis, while ‘*survival’*, ‘*lme4’* and ‘*MuMIn’* respectively for model fitting and averaging.

### Mapping grazing risk

We coupled the models describing the urchin use of the habitat and the kelp survival to create a map representing the risk for a kelp of being grazed in the different areas of the bottom. Using the ‘urchin use of the habitat’ model, we first mapped the urchin frequency assigning to each cell the maximum value between the maxima of predicted urchin presence probabilities in its parcels at different times. We then used this map as the input for the kelp survival model. The output of a CoxPH model is the log-hazard ratio, log(HR), here representing the risk for a kelp to be grazed and expressed relative to a cell with urchin frequency equal to 0. When the log(HR) equals 0 then two cells have the same risk.

## Results

### Site description

The spatial configuration of our study site was characterized by 2 sandy patches divided by one continuous patch of rocky bottom (Fig. 1b-c) however the two bottom types were almost equally represented (27:37, sandy:rocky cells).

Rocky patches were mainly composed by cobbles and gravel with boulder covering the 5.73 ± 0.1 % (mean ± standard error) while sand accounted for the 19.1± 3.0 % (mean ± s.e.). Sandy patch composition was more homogeneous with sand covering the 91.3 ± 1.7 % of the bottom, yet CAG and BLD were sparsely represented (7.0 ± 1.6 % and 1.5 ± 0.5 % mean ± s.e. respectively). Throughout the experiment the density of visible urchins on the rocky patch was of 0.92 ± 0.08 ind. m^-2^ (mean ± standard error, N=740) and of 0.13 ± 0.02 ind. m^-2^ (mean ± standard error, N=540) on the sandy patches.

### Kelp survival

Only 40 % of initially outplanted kelp survived at the end of the study (Fig. 2). We observed a sharp decrease in survival (40 % decrease) in the first 25 days followed by a constant decrease. Kelp survival was significantly related to the local urchin frequency (log HR ± .95 c.i. = 5.8 ± 2.9; LR: χ^2^ = 21.14, 1 df, P <0.001; table 1). The risk exposure decreased (i.e. Hazard ratio < 1) as urchin frequency decreased from the average value towards zero, while it greatly increased (Hazard ratio > 1) as urchin were more frequently present in a cell (Fig. 3).

**Figure 2.**
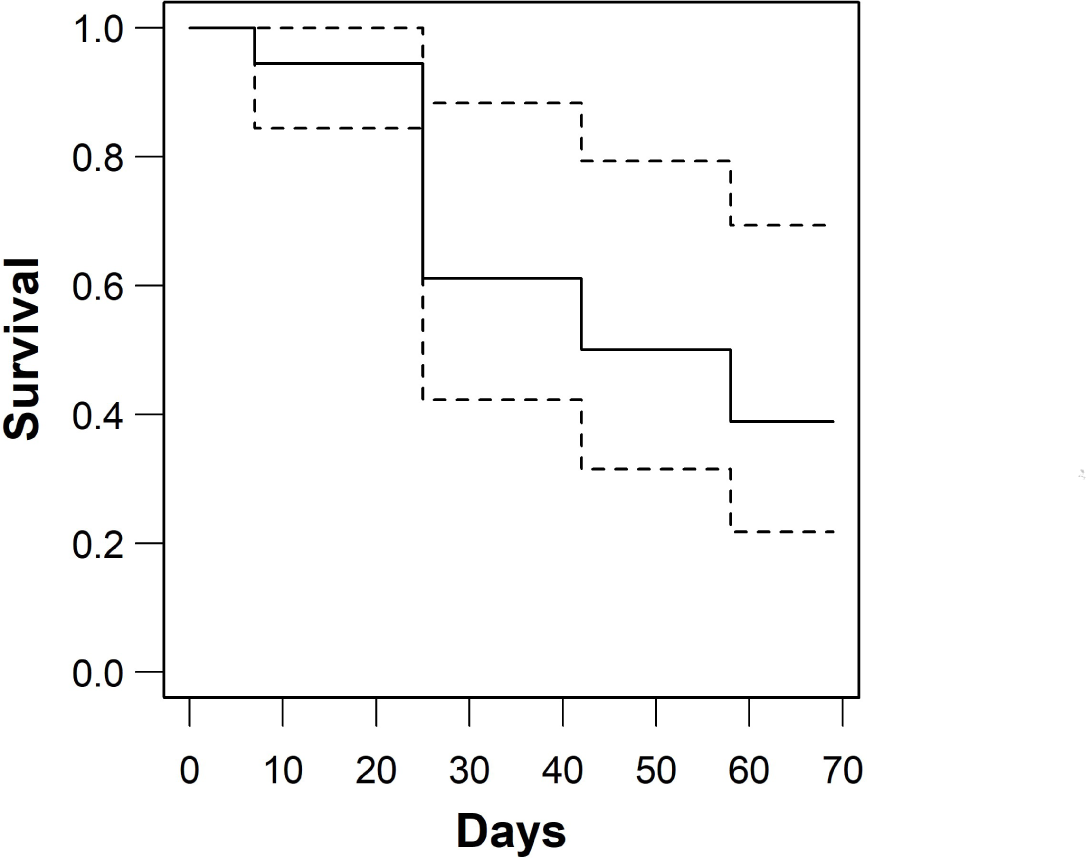
Kaplan-Maier survival curve of kelp. Vertical segments represent changes in survival at sampling times (x-axis, days from outplants). Solid line represent the kelp survival while dashed lines represent the 95 % confidence intervals.

**Figure 3.**
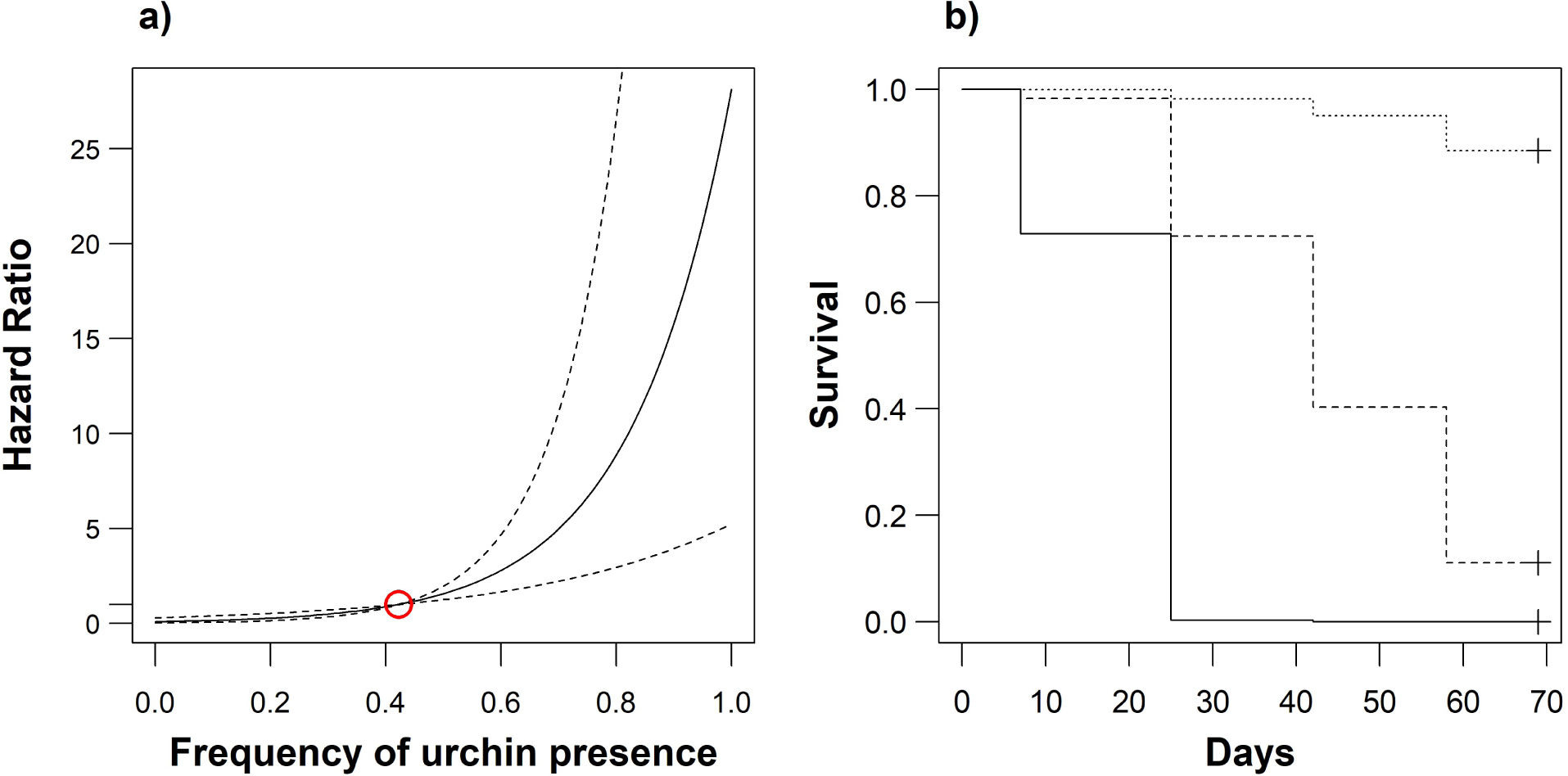
Fitted Cox Proportional Hazard model for kelp survival. a) Hazard ratio as function of urchin frequency. The hazard ratio is calculated relatively to the average urchin frequency, i.e. 0.42. dashed lines show the 0.95 confidence interval b) Predicted survival curves for 3 levels of urchin presence (0: dotted line, 0.5: dashed line, 1: solid line).

### Urchin use of the habitat

Three models resulted as good candidates between the ones tested and were therefore averaged (Table 2). The averaged model retained the variables BLD, CAG, Algae, CCA, AlgaeNeib and dBLOCK. All variables, except dBLOCK, were positively correlated with urchin frequency, although only BLD and CAG were significant and CCA was only almost significant (Table 3). The percentage cover of boulder in a parcel (BLD) had the greater effect size, meaning that an increase in boulder cover contribute the most in the frequency of urchin presence (Fig. 4). The averaged model showed a good calibration (i.e. a good agreement between observed outcomes and predictions) for predicted values below 0.4, otherwise the model seemed to slightly underestimate the observed values (Fig. S4). Model discrimination (i.e. the ability of model predicted values to discriminate between those with and without the outcome), assessed by the area under the receiver curve was satisfactory (AUC ± .95 c.i. = 0.95 ± 0.01; perfect discrimination when AUC is equal 1).

**Table 3.** Averaged model summary. Coefficients are shrinked estimates resulting from the ‘full’ average (i.e. considering a parameter equal to zero when not present in a given model). Values are in the linear predictors scale.

|  | Estimate | Adjusted SE | z value | P value |
| --- | --- | --- | --- | --- |
| Intercept | -5.406 | 0.604 | 8.96 | < 0.001 |
| BLD | 4.681 | 0.973 | 4.81 | < 0.001 |
| CAG | 2.191 | 0.502 | 4.36 | < 0.001 |
| Algae | 0.882 | 0.886 | 1.00 | 0.32 |
| CCA | 2.178 | 1.090 | 2.00 | 0.05 |
| AlgaeNeib | 1.562 | 1.097 | 1.42 | 0.15 |
| dBLOCK | -0.205 | 0.195 | 1.05 | 0.29 |

**Figure 4.**
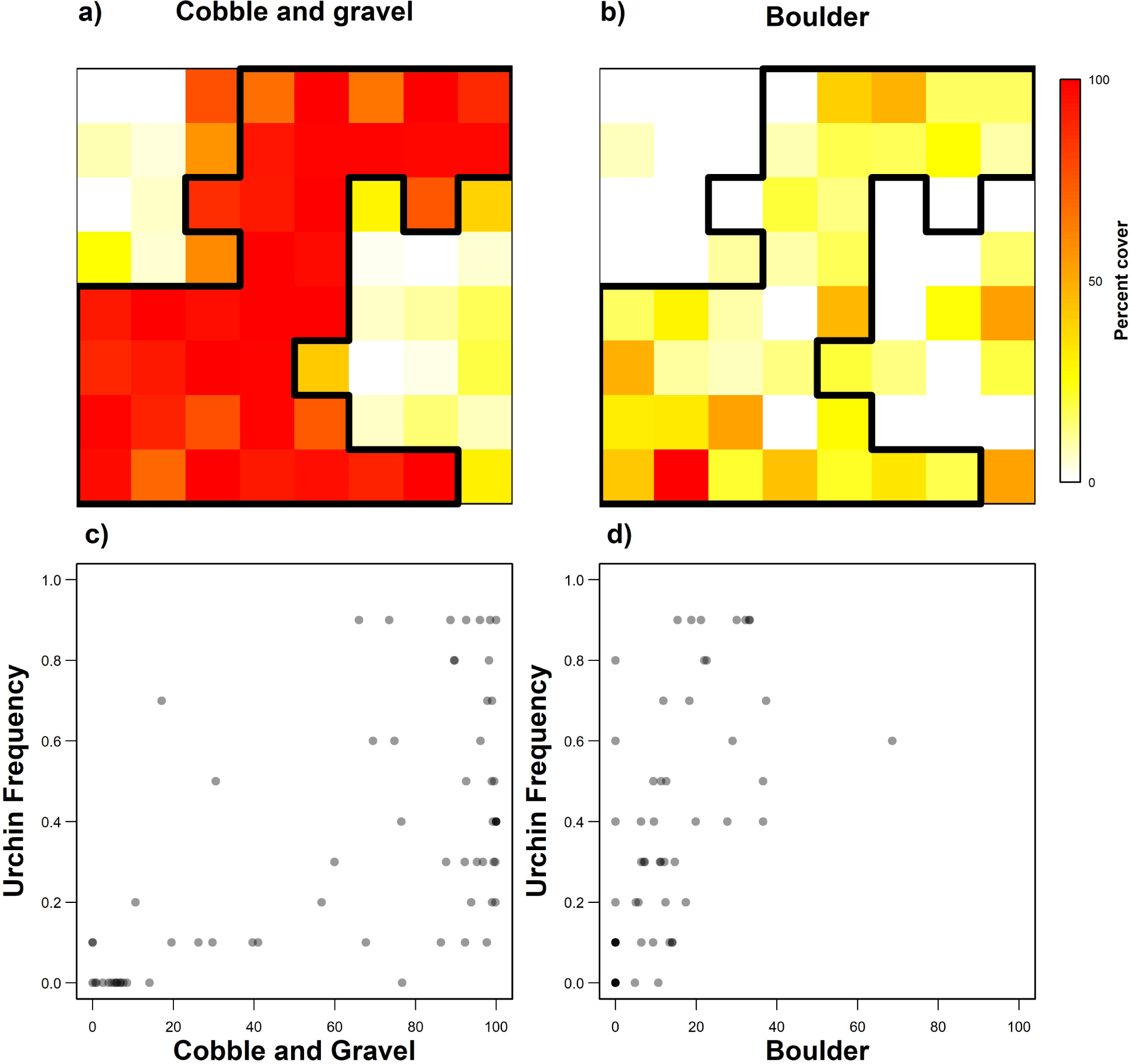
Distribution of gravel and boulders and their effect on urchin frequency. Heatmaps of a) cobble and gravel and b) boulders percent cover and relationship with the observed urchin frequency per cell in scatterplots c) and d) respectively. The black outline delimits the border of the rocky patch.

### Map of grazing risk

The grazing risk was concentrated on, but not limited to the rocky patch where it was generally higher than on the sandy part of the bottom (Fig. 5). The distribution of risk intensity also differed in the two patch types: risk intensity in rocky cells ranged between 0.6-5.2 and presented an overall uniform distribution, while log hazard ratio on sand ranged between 0-4.0 but 55 % of the cells were associated with the lowest risk (Fig. S5).

**Figure 5.**
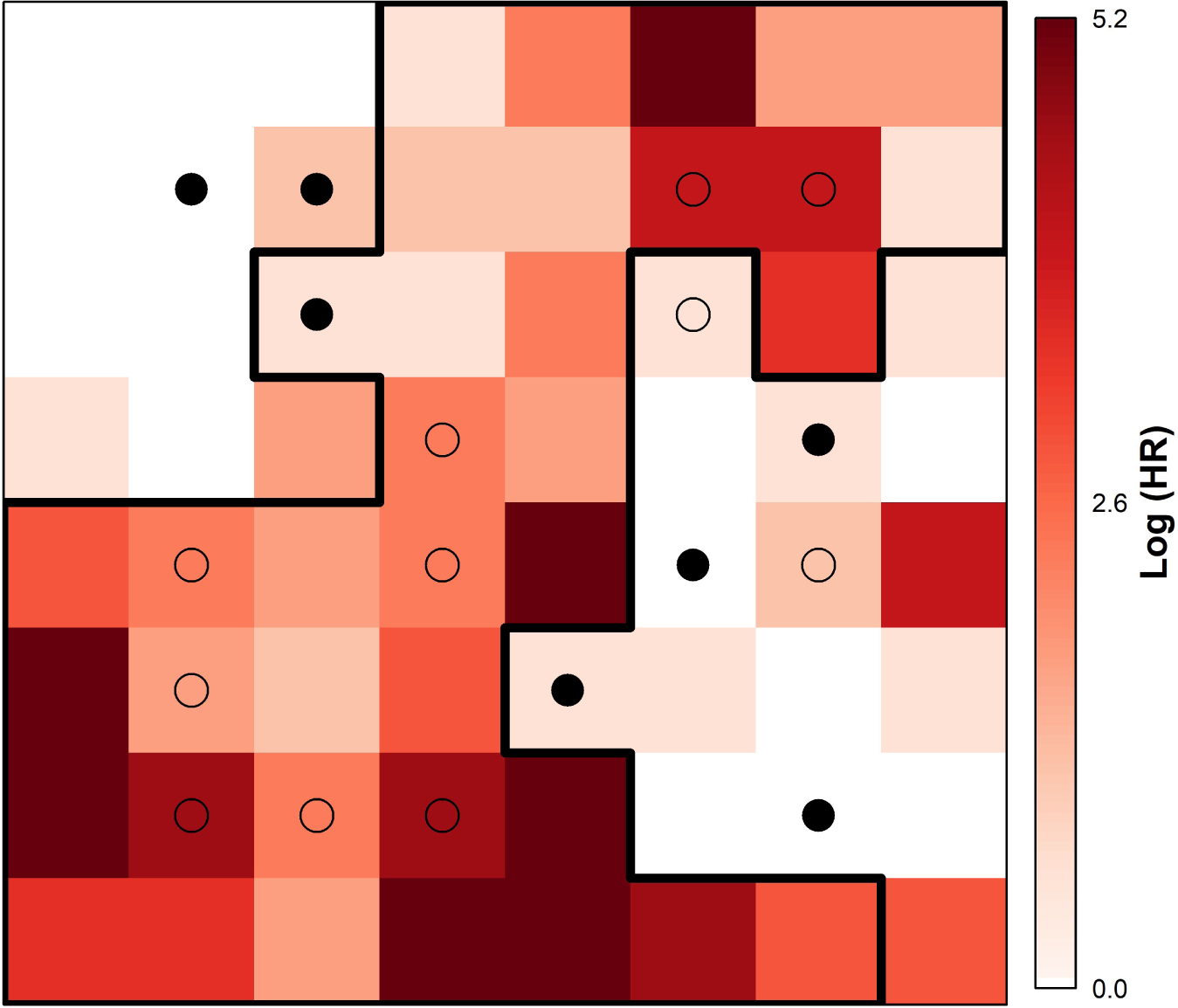
Grazing risk map. The risk is expressed as log hazard ratio (log(HR)) relative to cell with urchin frequency equal to 0. The darker the color the higher the risk of a kelp of being grazed; white cells have log(HR) equal zero and represent the reference level. The black outline delimits the border of the rocky patch. Black dots depict the position of survived kelp bot at the end of the experiment and one year after, open circles represent lost kelp.

A visual analysis of the map of grazing risk revealed the existence of hot-spots of risk in both rocky and sandy regions (i.e. darker cells in Fig. 5): cells associated with the highest risk were distributed at different location in the grid and separated from each other by lower risk cells. The position of the risk hot-spots seemed to be driven primarily by the distribution of boulders in the grid. Indeed, figure 4 showed that an increase in the percent cover of boulders in a given cell determined a more direct increase of urchin frequency compared to the amount of cobbles and gravel (CAG, the other significant variable in the model, Table 3).

Interestingly, survived kelp at the end of the experiment in 2015 were still alive one year after (Fig. 5).

### Herbivory exclusion and transplant mortality

In the absence of urchin herbivores (i.e. on the kelp frames) 83 % of kelp transplants survived and all developed reproductive sporophylls (sporophyll length = 26.1 ± 0.1 cm mean ± s.e., with no significant effect of depth, Fig. S6), indicating that very little mortality was due to the transplant technique and that the physical environment was suitable for kelp growth. There was no statistical effect of depth on the number of lost transplants (P=0.17, GLM with a Poisson distribution).

## Discussion

Using a spatially explicit framework, we investigated for the first time the role of the local seascape heterogeneity in modulating the interaction between the green sea urchin *S. droebachiensis* and kelp. We showed that grazing risk intensity is modulated by the seascape arrangement and its biological features, mainly bottom composition and algal cover, so that the presence of discrete seascape features can locally increase the grazing risk for kelp by differentially affecting the herbivore’s usage of the habitat, even within the same bottom patch. Importantly, we showed that even at a small spatial scale of tens of meters we can predict areas of lower grazing risk, i.e. kelp refugia, based on local seascape heterogeneity. While both the kelp-urchin dynamic (e.g. Ling et al., 2015; Robert E Scheibling, Hennigar, & Balch, 2011; Steneck et al., 2002) and the design and functioning of artificial reefs (Bulleri & Chapman, 2010; Fabi et al., 2011; Feary, Burt, & Bartholomew, 2011; Lima, Zalmon, & Love, 2019, and refeferences therein) have been frequently addressed in the literature, most of these studies are usually structured with a binary mindset that contrast alternative habitats, i.e. kelp bed vs. barren ground or artificial vs. natural reefs. By specifically considering the seascape heterogeneity, we offer a novel seascape framework within which artificial reef planning can take into consideration structuring ecological processes.

Seascape heterogeneity often results from multiple concurrent organizational levels (Didham, 2010). In this study, we recognized and specifically targeted heterogeneity at three levels: the bottom patchiness, the within-patch variability and the spatial configuration of features.

The patchiness of bottom composition played a major role in determining kelp refugia. Sandy patches were overall safer for kelp compared to rocky areas (Fig. 5). *S. droebacheinsis* prefer hard substrata, so this was an expected outcome. However, it confirmed a general predictive value of the coarse bottom classification into binary categories, i.e. sand vs rock, for the localization of kelp refugia in the seascape.

We detected a more interesting effect of the within-patch heterogeneity on the variability of grazing risk. Indeed, the intensity of grazing risk did not follow an all-or-nothing pattern matching the binary bottom classification, rather it was characterized by a higher within-patch variability with the presence of risk “hot-spots” (Fig. 5). Thanks to our two-step experimental approach coupling kelp survival with sea urchin habitat use, we resolved patch heterogeneity at a finer scale identifying sub-components affecting kelp survival. Specifically, the presence and distribution of boulders emerged as the primary seascape feature that determined the occurrence of hotspots of grazing risk both on rocky and sandy patches. Scheibling and Hamm (1991) reported that sea urchins greatly favored clusters of boulders over an adjacent cobble bed, since these microhabitats offered shelter form predation. On more uniform bedrock bottoms, *S. droebachiensis* also prefers microhabitats that favor anchorage, such as crevices, troughs and boulders, in the attempt to minimize dislodgement (Frey & Gagnon, 2016). Whether to avoid predation or dislodgement, structurally complex microhabitats drove the urchin distribution at our study site on the rocky matrix but also on sandy bottoms by locally enhancing the heterogeneity.

The cover of foliose and coralline algae contributed to the within-patch heterogeneity, particularly over the rocky bottom, increasing the urchin frequency. Foliose algae are a potential food source for urchins and, as such, their presence would either attract or retain urchin on the spot. Although we did not identify foliose algae to the species level, common red algae in the Saint-Lawrence Gulf are *Ptilota serrata* and *Phycodrys rubens* which have grazing-resistant properties (Himmelman, 1991). Thus alternatively, red foliose algae could limit the movement of urchins by offering an unpalatable obstacle or by physically disturbing the urchins (B & JA, 2003; Patrick Gagnon, Louis Vincent St-Hilaire-Gravel, Himmelman, & Johnson, 2006). In both cases, *S. droebachiensis* moving randomly in an area covered by foliose algae (C P Dumont, Himmelman, & Robinson, 2007; Lauzon-Guay, Scheibling, & Barbeau, 2006), could more frequently encounter algal patches increasing the chances to visit the same location multiple times. Regardless of the mechanisms involved, the percent cover of foliose red algae both in focal and neighboring regions had a predictive value for urchin presence and therefore for grazing risk in the area. Similarly, the percent cover of encrusting coralline algae is to be interpreted as a predictor of urchin presence, more likely the result of urchin grazing activity rather than a resource.

Both between-species and species-seascape interactions can be considered as an emergent property of the heterogeneity due to the spatial configuration of the landscape (Didham, 2010). For example, meadow proximity influences the trajectory of free-ranging bison moving in a patchy forest landscape (Dancose, Fortin, & Xulin, 2011), corridors of vegetation showing low-resistance to movement ease the spread of mountain pine beetles (Powell et al., 2018), while the spatial arrangement of seagrass affected the predation on sea urchins (Farina et al., 2016). Our models included variables describing the spatial configuration of both the seascape and the artificial blocks (Table 2), however the limited spatial extent of our study or the abundance of cells containing boulders (Fig. 4) possibly hindered our ability to capture an effect of seascape configuration. Nonetheless, the slightly negative effect of the distance from a block on urchin frequency suggests that blocks could possibly act as a sink. Indeed, we commonly observed urchins occupying the block cavities (Fig. S7), a behavior consistent with the urchin’s preference for boulder’ sheltered microhabitats (R. E. Scheibling & Hamm, 1991, and this study).

Grazing risk for kelp is thus an emerging property resulting from the interaction between the green sea urchin and the local habitat. The survival of kelp was positively correlated to the frequency of urchin presence in a given cell, pointing at urchin movement throughout the seascape as the underlying mechanism. Individual movement ultimately determine the distribution of urchin density, which in our case resulted the second-best explanatory variable (Table 1). Drastic changes in kelp cover are known to happen with urchin density in the order of tens per meter squared (Filbee-Dexter & Scheibling, 2014), although densities as low as 4 ind. m^-2^ have been recorded in transition areas between kelp bed and urchin barren (Jeon, Yang, & Kim, 2015). In our study, grazing impact even at low densities support the perspective that individual movement could become increasingly important as urchin density decrease (Lauzon-Guay et al., 2006). Tracking the movement of individual urchins and relating it to the seascape would have certainly been the ideal approach for our study, but unfortunately is extremely complex and demanding. However, by using the frequency of urchin presence in a cell as a proxy for movement, we can hypothesize a mechanism underlying the emergence of differential grazing risk areas. The preference of *S. droebachiensis* for structurally complex features, such as boulders, determine areas of increased risk over the overall high-risk rocky bottom patches, the typical substrate for urchin where they move randomly. Generally safer, sandy bottoms do not necessarily represent an obstacle to urchin movement so that some individuals venture on soft substrate and can find artificial blocks. How *S. droebachiensis* moves on sand is still not known but could be at the base of low grazing risk of kelp on soft bottom. Dumont et al. (2006) observed that *S. droebachiensis* could move further on sand than they would on a rocky bottom. Adopting a “sand-movement mode” (e.g. longer, straighter trajectories), *S. droebachiensis* could minimize permanence on the sand to avoid predation (Levitan & Genovese, 1989), decreasing the chance of finding artificial blocks and kelp blades lying on the bottom. If found, we hypothesize that the survival of these plants could depend on the blade escaping from the urchin grab, once shortened, thus remaining in a more elevated position: *Alaria* stipes were in fact anchored on top of blocks. Urchin sheltering in the block cavities will have more chances to step foot on sand rather than making it to the top of the block. This would potentially trigger the “sand-movement mode” resulting in the urchin more easily leaving the area rather than finding the block a second time, and thus in a lower grazing risk for the kelp on the block.

Seascape ecology offers a conceptual, research and applied framework to identify areas with of differential intensity of species interaction. In this sense, our work offers an example of the predictive potential of seascape features to minimize grazing risk, even at the small scale at which artificial reefs are used in restoration and habitat enhancement projects. Such a predictive capability opens to the possibility to better integrate artificial reefs in a more coherent ecological context. Deploying artificial reef modules closer to natural rocky bottoms could help reducing some drawbacks and impacts of past approaches to minimize grazing on artificial structures, while promoting desirable aspects of connectivity between artificial and natural habitats (Bishop et al., 2017). For example, to deter urchins from reefs, modules have been placed over increasingly, when not isolated on, sandy bottom areas, or their height have been raised to the depth of hard wave motion (Reed, Schroeter, Huang, Anderson, & Ambrose, 2006; T Terawaki, Arai, & Kawasaki, 1995; Toshinobu Terawaki, Hasegawa, Arai, & Ohno, 2001; personal observations). Increasing a structure’s height implies higher costs to deploy more material and to assure structural stability. At high latitudes kelp persistence could also be reduced by exposing to ice scouring the grazing-safer, shallower parts of a structure. Ecologically, the introduction of extraneous hard substrate on soft sediments alters the existing habitat affecting the local hydrodynamic and the composition of sediments, augments organic matter inputs, and increases predatory and grazing pressure on the surrounding soft bottom communities generating halos (Bishop et al., 2017; Heery et al., 2017). Grazers could still reach isolated structures, either through recruitment or movement, clearing the artificial macroalgal bed and creating opportunities for invasive species (Dafforn, Glasby, & Johnston, 2012). On the contrary, arranging kelp-covered reefs modules within reduced grazing risk areas in proximity of existing rocky bottoms could promote positive feedback on natural habitats. For example, a recent survey of an isolated modular reef in the Bay of Sept-Iles, Canada, estimated an average of 3656 gm^-2^ of kelp biomass at the artificial reef site scale, mainly adult fronds, with up to 14200 gm^-2^ on average per reef unit (Ferrario & Archambault, 2018). If similar yields were to be close to natural rocky reefs, they would represent an important contribute to the system. Indeed, kelp fragments drifting from reef units onto barren ground would enhance feeding opportunities for local sea urchin populations, favoring the development of their gonads and enhancing their commercial value (Claisse et al., 2013; Cresson, Ruitton, & Harmelin-Vivien, 2014). Spore supplies could support the recruitment of juvenile sporophytes on nearby rocks, creating the premises for the growth of natural kelp bed when favorable condition may arise.

Our modelling approach offers a method to inform stakeholder with both a prediction of where urchin grazing risk will be lowest at the site scale and an indication on which predictor variables they need to survey. We thus believe that the usefulness of our seascape approach is greatest particularly when it comes to guide restoration projects since these are necessarily implemented at the local scale, where ultimately their outcome depends on the intensity of species interactions. On the contrary, seascape approaches and species distribution models that studied the distribution of kelp beds and urchin barren have been so far conducted at large scales and only using abiotic environmental parameters as predictors (e.g. Parnell, 2015; Rinde et al., 2014). While we recognize the relevance of these studies in highlighting general trends and spatial patterns, their usefulness weakens when trying to shed light on the outcomes of any species interaction and their underlying mechanisms.

Several factors need to be considered to promote a more widespread adoption of our approach. Primarily, the number of species involved in the interaction, their ecology and their degree of mobility needs to be clearly assessed. In our case, *S. droebachiensis* was the only relevant grazer in the system and well-studied species, allowing us to define a suitable scale of experimentation and the set of variables to be mapped. In particular, the degree of species mobility and the ecology of the species determine the spatial scale and resolution at which seascape data need to be acquired. Unfortunately, this information is virtually absent in the majority of coastal context. We thus encourage environmental managers, regional and local governmental bodies to urge the mapping of coastal habitats, with a particular attention to biological data (e.g. vegetation and benthic invertebrates). Reports on artificial reefs projects are frequently kept private and/or projects outcome are not monitored (Tessier et al., 2015). This information needs to be made publicly available to ease the identification of potential factors affecting the biotic disturbance (e.g. grazing).

Our work demonstrates that we can understand how the intensity of species interactions is modulated by the surrounding seascape by explicitly considering the spatial heterogeneity even at local scales. This information can be readily incorporated in the planning phase of modular artificial reefs to minimize the exposition of individual reef units to detrimental grazing risk thus increasing their rate of success and ensuring lasting results.

## Authors’ contributions

FF and LEJ conceived the idea and designed the experiment; FF and TS performed the experiment; FF analyzed the data; FF led the writing of the manuscript. All authors contributed critically to the drafts and gave final approval for publication.

## References

Airoldi, L., & Bulleri, F. (2011). Anthropogenic disturbance can determine the magnitude of opportunistic species responses on marine urban infrastructures. PLoS ONE, 6(8), e22985. Retrieved from http://dx.doi.org/10.1371%2Fjournal.pone.0022985

Airoldi, L., Connell, S. D., & Beck, M. W. (2009). The loss of natural habitats and the addition of artificial substrata. In M. Wahl (Ed.), Marine Hard Bottom Communities: Patterns, Dynamics, Diversity, and Change (Vol. 206, pp. 269–280). doi: 10.1007/978-3-540-92704-4_19

B, K., & Ja, E. (2003). The stability of boundary regions between kelp beds and deforested areas. Ecology, 84(1), 174. doi: 10.1890/0012-9658(2003)084[0174:tsobrb]2.0.co;2

Bishop, M. J., Mayer-Pinto, M., Airoldi, L., Firth, L. B., Morris, R. L., Loke, L. H. L., … Dafforn, K. A. (2017). Effects of ocean sprawl on ecological connectivity: impacts and solutions. Journal of Experimental Marine Biology and Ecology, 492, 7–30. doi: 10.1016/j.jembe.2017.01.021

Boström, C., Pittman, S. J., Simenstad, C., & Kneib, R. T. (2011). Seascape ecology of coastal biogenic habitats: Advances, gaps, and challenges. Marine Ecology Progress Series, 427, 191–217. doi: 10.3354/meps09051

Bulleri, F., & Chapman, M. G. (2010). The introduction of coastal infrastructure as a driver of change in marine environments. Journal of Applied Ecology, 47(1), 26–35. doi: 10.1111/j.1365-2664.2009.01751.x

Carney, L. T., Waaland, J. R., Klinger, T., & Ewing, K. (2005). Restoration of the bull kelp Nereocystis luetkeana in nearshore rocky habitats. Marine Ecology Progress Series, 302, 49–61. doi: 10.3354/meps302049

Claisse, J. T., Williams, J. P., Ford, T., Pondella, D. J., Meux, B., & Protopapadakis, L. (2013). Kelp forest habitat restoration has the potential to increase sea urchin gonad biomass. Ecosphere, 4(3), art38. doi: 10.1890/ES12-00408.1

Cresson, P., Ruitton, S., & Harmelin-Vivien, M. (2014). Artificial reefs do increase secondary biomass production: Mechanisms evidenced by stable isotopes. Marine Ecology Progress Series, 509, 15–26. doi: 10.3354/meps10866

Dafforn, K. A., Glasby, T. M., & Johnston, E. L. (2012). Comparing the invasibility of experimental “reefs” with field observations of natural reefs and artificial structures. PLoS ONE, 7(5), e38124. doi: e38124 10.1371/journal.pone.0038124

Dancose, K., Fortin, D., & Xulin, G. U. O. (2011). Mechanisms of functional connectivity: The case of free-ranging bison in a forest landscape. Ecological Applications, 21(5), 1871–1885. doi: 10.1890/10-0779.1

Didham, R. K. (2010). Ecological Consequences of Habitat Fragmentation. In Encyclopedia of Life Sciences. doi: 10.1002/9780470015902.a0021904

Dumont, C P, Himmelman, J. H., & Robinson, S. M. C. (2007). Random movement pattern of the sea urchin Strongylocentrotus droebachiensis. Journal of Experimental Marine Biology and Ecology, 340(1), 80–89. doi: 10.1016/j.jembe.2006.08.013

Dumont, Clément P., Himmelman, J. H., & Russell, M. P. (2006). Daily movement of the sea urchin Strongylocentrotus droebachiensis in different subtidal habitats in eastern Canada. Marine Ecology Progress Series, 317, 87–99. doi: 10.3354/meps317087

Dyson, K., & Yocom, K. (2015). Ecological design for urban waterfronts. Urban Ecosystems, 18(1), 189–208. doi: 10.1007/s11252-014-0385-9

Fabi, G., Spagnolo, A., Bellan-Santini, D., Charbonnel, E., Cicek, B. A., Garcia, J. J. G., … dos Santos, M. N. (2011). OVERVIEW ON ARTIFICIAL REEFS IN EUROPE. Brazilian Journal of Oceanography, 59, 155–166.

Farina, S., Guala, I., Oliva, S., Piazzi, L., Pires da Silva, R., & Ceccherelli, G. (2016). The Seagrass Effect Turned Upside Down Changes the Prospective of Sea Urchin Survival and Landscape Implications. PLOS ONE, 11(10), e0164294. doi: 10.1371/journal.pone.0164294

Feary, D. A., Burt, J. A., & Bartholomew, A. (2011). Artificial marine habitats in the Arabian Gulf: Review of current use, benefits and management implications. Ocean and Coastal Management, 54(10), 742–749. doi: 10.1016/j.ocecoaman.2011.07.008

Ferrario, F., & Archambault, P. (2018). Portrait de la communauté de macroalgues de la zone subtidale. In J. Carrière (Ed.), Observatoire environnemental de la baie de Sept-Îles (pp. 491–566). Retrieved from http://www.inrest.ca/environnement/projets?real=1

Ferrario, F., Iveša, L., Jaklin, A., Perkol-Finkel, S., & Airoldi, L. (2016). The overlooked role of biotic factors in controlling the ecological performance of artificial marine habitats. Journal of Applied Ecology, 53(1), 16–24. doi: 10.1111/1365-2664.12533

Filbee-Dexter, K., & Scheibling, R. E. (2014). Sea urchin barrens as alternative stable states of collapsed kelp ecosystems. Marine Ecology Progress Series, 495, 1–25. doi: 10.3354/meps10573 *Fisheries Act*., (2019).

Fortin, D., Beyer, H. L., Boyce, M. S., Smith, D. W., Duchesne, T., & Mao, J. S. (2005). Wolves influence elk movements: behavior shapes a trophic cascade in yellowstone national park. Ecology, 86(5), 1320–1330. doi: 10.1890/04-0953

Foster, M. S. (1990). Organization of macroalgal assemblages in the Northeast Pacific: the assumption of homogeneity and the illusion of generality. Hydrobiologia, 192(1), 21–33. doi: 10.1007/BF00006225

Frey, D. L., & Gagnon, P. (2016). Spatial dynamics of the green sea urchin Strongylocentrotus droebachiensis in food-depleted habitats. Marine Ecology Progress Series, 552, 223–240. doi: 10.3354/meps11787

Gagnon, P., Himmelman, J. H., & Johnson, L. E. (2004). Temporal variation in community interfaces: Kelp-bed boundary dynamics adjacent to persistent urchin barrens. Marine Biology, 144(6), 1191–1203. doi: 10.1007/s00227-003-1270-x

Gagnon, Patrick, Louis Vincent St-Hilaire-Gravel, Himmelman, J. H., & Johnson, L. E. (2006). Organismal defenses versus environmentally mediated protection from herbivores: Unraveling the puzzling case of Desmarestia viridis (Phaeophyta). Journal of Experimental Marine Biology and Ecology, 334(1), 10–19. doi: 10.1016/j.jembe.2006.01.012

Gil, M. A., Zill, J., & Ponciano, J. M. (2017). Context-dependent landscape of fear: algal density elicits risky herbivory in a coral reef. Ecology, 98(2), 534–544. doi: 10.1002/ecy.1668

Heery, E. C., Bishop, M. J., Critchley, L. P., Bugnot, A. B., Airoldi, L., Mayer-Pinto, M., … Dafforn, A. (2017). Identifying the consequences of ocean sprawl for sedimentary habitats. Journal of Experimental Marine Biology and Ecology, 492, 31–48. doi: 10.1016/j.jembe.2017.01.020

Himmelman, J. H. (1991). Diving observation of subtidal communities in the northern Gulf of Saint Lawrence. Canadian Special Publication of Fisheries and Aquatic Sciences, 113(113), 319–332.

Jeon, B. H., Yang, K. M., & Kim, J. H. (2015). Changes in macroalgal assemblage with sea urchin density on the east coast of South Korea. Algae, 30(2), 139–146. doi: 10.4490/algae.2015.30.2.139

Keats, D. W. (1991). Refugial Laminaria Abundance and Reduction in Urchin Grazing in Communities in the North-West Atlantic. Journal of the Marine Biological Association of the United Kingdom, 71(4), 867–876. doi: 10.1017/s0025315400053510

Kleinbaum, D. G., & Klein, M. (2005). Survival analysis a self-learning text. Retrieved from http://ariane.ulaval.ca/cgi-bin/recherche.cgi?qu=i9780387291505

Lauzon-Guay, J. S., Scheibling, R. E., & Barbeau, M. A. (2006). Movement patterns in the green sea urchin, Strongylocentrotus droebachiensis. Journal of the Marine Biological Association of the United Kingdom, 86(1), 167–174. doi: 10.1017/S0025315406012999

Levitan, D. R., & Genovese, S. J. (1989). Substratum-dependent predator-prey dynamics: patch reefs as refuges from gastropod predation. Journal of Experimental Marine Biology and Ecology, 130(2), 111–118. doi: 10.1016/0022-0981(89)90198-6

Lima, J. S., Zalmon, I. R., & Love, M. (2019). Overview and trends of ecological and socioeconomic research on artificial reefs. Marine Environmental Research, 145, 81–96. doi: 10.1016/j.marenvres.2019.01.010

Ling, S. D., Scheibling, R. E., Rassweiler, A., Johnson, C. R., Shears, N., Connell, S. D., … Johnson, E. (2015). Global regime shift dynamics of catastrophic sea urchin overgrazing. Philosophical Transactions of the Royal Society B: Biological Sciences, 370(1659), 1–10. doi: 10.1098/rstb.2013.0269

Madin, E. M. P., Madin, J. S., & Booth, D. J. (2011). Landscape of fear visible from space. Scientific Reports, 1, 4. doi: 1410.1038/srep00014

Mann, K. H. (1973). SEAWEEDS - THEIR PRODUCTIVITY AND STRATEGY FOR GROWTH. Science, 182(4116), 975–981. doi: 10.1126/science.182.4116.975

Mason, T. H. E., & Fortin, D. (2017). Functional responses in animal movement explain spatial heterogeneity in animal-habitat relationships. Journal of Animal Ecology, 86(4), 960–971. doi: 10.1111/1365-2656.12682

Mayer-Pinto, M., Johnston, E. L., Bugnot, A. B., Glasby, T. M., Airoldi, L., Mitchell, A., & Dafforn, K. A. (2017). Building “blue”: An eco-engineering framework for foreshore developments. Journal of Environmental Management, 189, 109–114. doi: 10.1016/j.jenvman.2016.12.039

Mazerolle, M. J. (2006). Improving data analysis in herpetology: using Akaike’s Information Criterion (AIC) to assess the strength of biological hypotheses. Amphibia-Reptilia, 27(2), 169–180. doi: 10.1163/156853806777239922

Micheli, F., & Peterson, C. H. (1999). Estuarine vegetated habitats as corridors for predator movements. Conservation Biology, 13(4), 869–881. doi: 10.1046/j.1523-1739.1999.98233.x

Parnell, P. E. (2015). The effects of seascape pattern on algal patch structure, sea urchin barrens, and ecological processes. Journal of Experimental Marine Biology and Ecology, 465(0), 64–76. doi: 10.1016/j.jembe.2015.01.010

Perkol-Finkel, S., Ferrario, F., Nicotera, V., & Airoldi, L. (2012). Conservation challenges in urban seascapes: promoting the growth of threatened species on coastal infrastructures. Journal of Applied Ecology, 49(6), 1457–1466. doi: 10.1111/j.1365-2664.2012.02204.x

Perkol-Finkel, S., Hadary, T., Rella, A., Shirazi, R., & Sella, I. (2018). Seascape architecture –incorporating ecological considerations in design of coastal and marine infrastructure. Ecological Engineering, 120, 645–654. doi: 10.1016/j.ecoleng.2017.06.051

Powell, J. A., Garlick, M. J., Bentz, B. J., & Friedenberg, N. (2018). Differential dispersal and the Allee effect create power-law behaviour: Distribution of spot infestations during mountain pine beetle outbreaks. Journal of Animal Ecology, 87(1), 73–86. doi: 10.1111/1365-2656.12700

R Core Team. (2016). R: a language and environment for statistical computing. Retrieved from http://www.r-project.org

Reed, D. C., Schroeter, S. C., Huang, D., Anderson, T. W., & Ambrose, R. F. (2006). Quantitative assessment of different artificial reef designs in mitigating losses to kelp forest fishes. Bulletin of Marine Science, 78(1), 133–150.

Rinde, E., Christie, H., Fagerli, C. W., Bekkby, T., Gundersen, H., Norderhaug, K. M., & Hjermann, D. (2014). The influence of physical factors on kelp and sea urchin distribution in previously and still grazed areas in the NE Atlantic. PLoS ONE, 9(6), e100222. doi: 10.1371/journal.pone.0100222

Scheibling, R. E., & Hamm, J. (1991). Interactions between sea urchins (Strongylocentrotus droebachiensis) and their predators in field and laboratory experiments. Marine Biology, 110(1), 105–116. doi: 10.1007/BF01313097

Scheibling, Robert E, Hennigar, A. W., & Balch, T. (2011). Destructive grazing, epiphytism, and disease: the dynamics of sea urchin - kelp interactions in Nova Scotia. Canadian Journal of Fisheries and Aquatic Sciences, 56(12), 2300–2314. doi: 10.1139/f99-163

Steneck, R. S., Graham, M. H., Bourque, B. J., Corbett, D., Erlandson, J. M., Estes, J. A., & Tegner, M. J. (2002). Kelp forest ecosystems: Biodiversity, stability, resilience and future. Environmental Conservation, 29(4), 436–459. doi: 10.1017/S0376892902000322

Steneck, R. S., Leland, A., McNaught, D. C., & Vavrinec, J. (2013). Ecosystem flips, locks, and feedbacks: The lasting effects of fisheries on Maine’s kelp forest ecosystem. Bulletin of Marine Science, 89(1), 31–55. doi: 10.5343/bms.2011.1148

Terawaki, T, Arai, S., & Kawasaki, Y. (1995). Methods of submarine forest formation considering local limiting factors of distribution (pp. 145–154; F. A. O. of the UN, Ed.). pp. 145–154.

Terawaki, Toshinobu, Hasegawa, H., Arai, S., & Ohno, M. (2001). Management-free techniques for restoration of Eisenia and Ecklonia beds along the central Pacific coast of Japan. Journal of Applied Phycology, 13(1), 13–17. doi: 10.1023/A:1008135515037

Tessier, A., Francour, P., Charbonnel, E., Dalias, N., Bodilis, P., Seaman, W., & Lenfant, P. (2015). Assessment of French artificial reefs: due to limitations of research, trends may be misleading. Hydrobiologia, 753(1), 1–29. doi: 10.1007/s10750-015-2213-5

Zuur, A. F., Ieno, E. N., Walker, N. J., Saveliev, A. A., & Smith, G. M. (2009). Mixed effects models and extensions in ecology with R. In M. Gail, K. Krickeberg, J. M. Samet, A. Tsiatis, & W. Wong (Eds.), Statistics for Biology and Health. doi: 10.1007/978-0-387-87458-6_1

