## Supporting Information for "Spatially explicit modelling of kelp-grazer interactions: a seascape approach for effective habitat enhancement using artificial reefs"

### Method

Bottom mapping

We deployed a coordinate system allowing us to navigate through the grid and reference photographs (Fig. S1). In the grid, the x-axis coordinates consisted of four parallel transect lines (anchored to the bottom four meters apart) delimiting two contiguous rows of cells (one per each side of the transect, i.e. line 1 between rows 1-2, line 2 between rows 3-4 and so forth). The y-axis coordinates consisted of tags spaced every two meters along each transect (eight per transect, numbered 1 to 8 and with the same orientation) which identified the position of the center of each cell (Fig. S1). To acquire data from each cell of the grid, we took four orthogonal photographs of the bottom using a GoPro Hero3$^+$ camera mounted on a PVC frame whose base, a 0.8×0.8 m square, was rotated clockwise pivoting on the center of the cell by the diver, virtually subdividing the cell into four unique parcels (Fig. S2). This strategy improved the clarity of each parcel and allowed for better maneuverability of the frame underwater. The camera was set to take a burst of 10 pictures in 2 seconds (12MP –Wide mode) and only the best picture from each series was subsequently analyzed.

Two divers operated in parallel on each sided of the same transect maintaining visual contact and, depending on their experience, were able to complete one transect in about 20 minutes.

Before analysis, we preprocessed all pictures using the automatic white balance adjustment feature in GIMP 2.8 (www.gimp.org) to enhance image clarity.

*Substrate Type* - We defined three categories of substratum type based on the Wentworth scale (Wentworth, 1922): “boulders” (BLD, >25cm), “cobbles and gravel” (CAG, 0.2-25 cm) and “sand” (SND, < 0.2 cm). Substrate size was estimated using 5 cm markers on the frame base as reference, while soft substrate was classified as sand by divers. Substrate size refer to the longer axis. BLD and CAG were further divided in two subcategories since they were usually either encrusted with coralline algae (BLD+ / CAG+ CCA) or covered by a layer of algal turf and sediment (BLD+ / CAG+ TurfSed). Additionally, the percent cover of foliose algae (ALG) was quantified. We measured the percent cover of each category inside the PVC frame using the software Vidana 1.1 (marinespatialecologylab.org). Each cell and its four parcels were designated as either a sandy or a rocky patch following a cluster analysis based on a Euclidean distance matrix of the per cell mean values of BLD, CAG, SND and ALG.

*Algal cover* – The percent cover of foliose algae (‘Algae’, mainly red algae), turf and sediment (‘TurfSed’), and crustose coralline algae (CCA) was estimated using the computer vision algorithms implemented in CoralNet (www.coralnet.ucsd.edu; Beijbom et al., 2015). Percent cover was estimated for each parcel based on 50 points randomly overlayed onto each image. The identity of the feature under each point was classified according to a predefined labelset which included the three algal categories of interest, common benthic invertebrates (e.g. urchin, crab), a category for the frame and one for the bare cement block. Points which had a confidence threshold below 95% from the machine learning classifier CoralNet (www.coralnet.ucsc.edu) were visually examined and manually verified.

Kelp outplanting

We affixed kelp onto artificial concrete blocks by inserting the holdfast of each thallus through a small cut in a rubber strip that was fixed onto a 6mm polypropylene rope with cable ties. This method allowed the holdfast to be in contact with the rope, and has been used extensively to transplant kelp plants on kelp farms (Fig S3). Each rope was numbered, and tied onto previously deployed artificial blocks by divers in the field. We deemed such outplanting technique effective based on previous similar experiences (Carney, Waaland, Klinger, & Ewing, 2005) during which the holdfast of outplanted kelp was observed to adhere to the rope, as it did in our study.

Verification of outplant technique and suitable kelp growing conditions

The kelp frame was composed of three polypropylene lines (12mm diameters, 11.3m total length). Each line was anchored at one end to one of three 25kg blocks, and the three lines attached to a single subsurface (1m depth MLLW) float. The resulting structure was pyramidal in shape, rigid, and with each line at a 45º angle from the bottom and 120º from the other two lines. This configuration allowed us to attach kelps to the frame along a depth gradient using the same materials (i.e. a rubber strip above the hold fast attached via cable-ties to a polypropylene line) used to throughout the rest of our experiment. The diagonal orientation of each line in the water column prevented shading of kelps attached closer to the bottom by those shallower on the frame (fig. S3b).

Statistical Analysis

*Survival analysis* - In our dataset only right censoring occurred and mainly because of kelp surviving until the end of the experimental follow-up.

For CoxPH survival models we graphically compared the observed versus the expected survival curve plots of the selected semi-parametric model to check the proportional hazard assumption (Kleinbaum & Klein, 2005). Survival analysis was conducted with the R package '*survival*' (Therneau, 2019)

*Urchin use of the habitat* - We checked for collinearity between all potential predictors by means of pairplots and retained the variables that we deemed more relevant within pairs having a correlation ≤ 0.6. Given the binary nature of the response variable "urchin presence/absence", we used generalized regression models with a binomial distribution and logit link. Following Zuur et al. (2009), the random effect "parcel" was included after AIC comparison of sub-global models (Glb1, Glb2, Glb3 in Table 2 in the main paper) with and without the random effect. Sub-global models were used because of numerical instability when adding a random component to the global model containing all the predictors.

The final set of candidate models included the three sub-global models, one model with predictors of each seascape property alone (i.e. bottom composition, biological factors, spatial configuration and the reef spatial arrangement) plus one for biological properties without neighboring predictor, and 9 combinations of properties pairs, for a total of 17 model formulations (table 2). The dataset used contained 1017 data points (256 parcels × 4 times, excluding 7 non-complete cases).

We checked the adequacy of each averaged model by graphically inspecting the quantile-quantile and partial residual plots to detect departure from model assumption as well as the spline correlogram to detect spatial correlation (Zuur et al., 2009). To improve linearity, percent covers were arcsin transformed and the whole model selection process was repeated. Models were fitted using the generalized linear mixed-model function *glmer* in the R package '*lme4*' (Bates, Mächler, Bolker, & Walker, 2015) while the '*MuMIn*' package was used for model averaging (Barton, 2016). Spatial analysis were performed using the package '*raster*' (Hijmans, 2016).

**Figures**

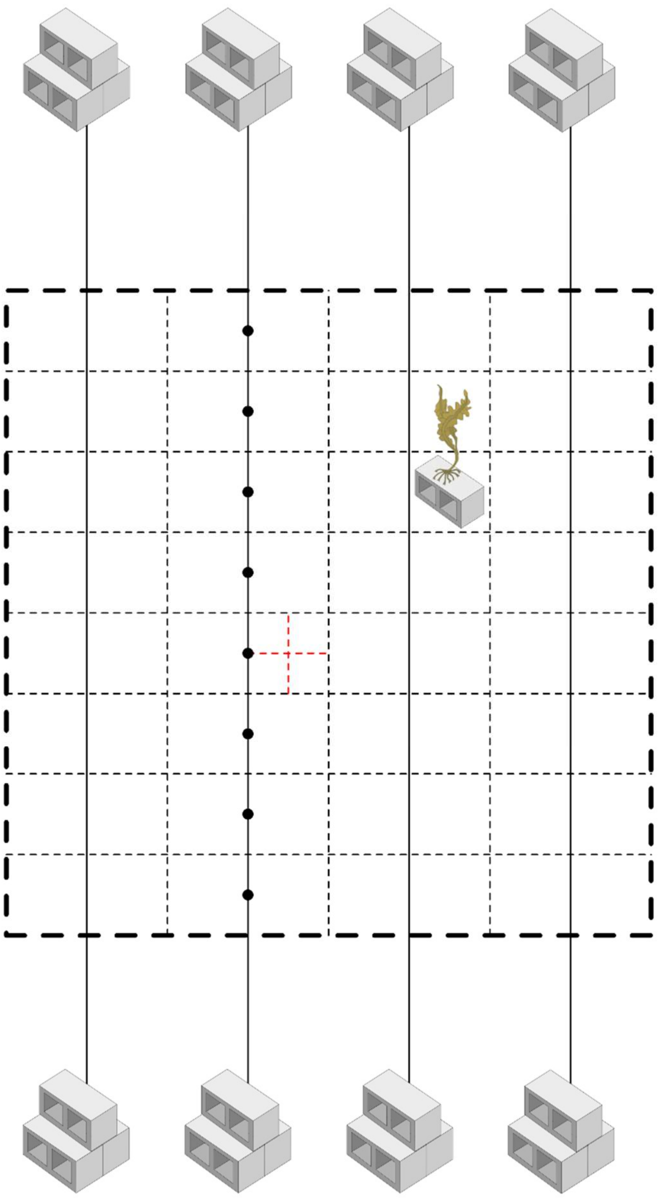

**Figure S1.** Schematic of the mapping grid. Four transect (solid line) were hold in position by cinder block anchors to identify the area to be mapped (dashed lines). Each cell of 2 meters side was virtually divided in four parcels (red dashed lines). Tags spaced along each transect referenced the position of the center of the cells. Graphic of the alga credit of Tracey Saxby, Integration and Application Network, University of Maryland Center for Environmental Science (ian.umces.edu/imagelibrary/).

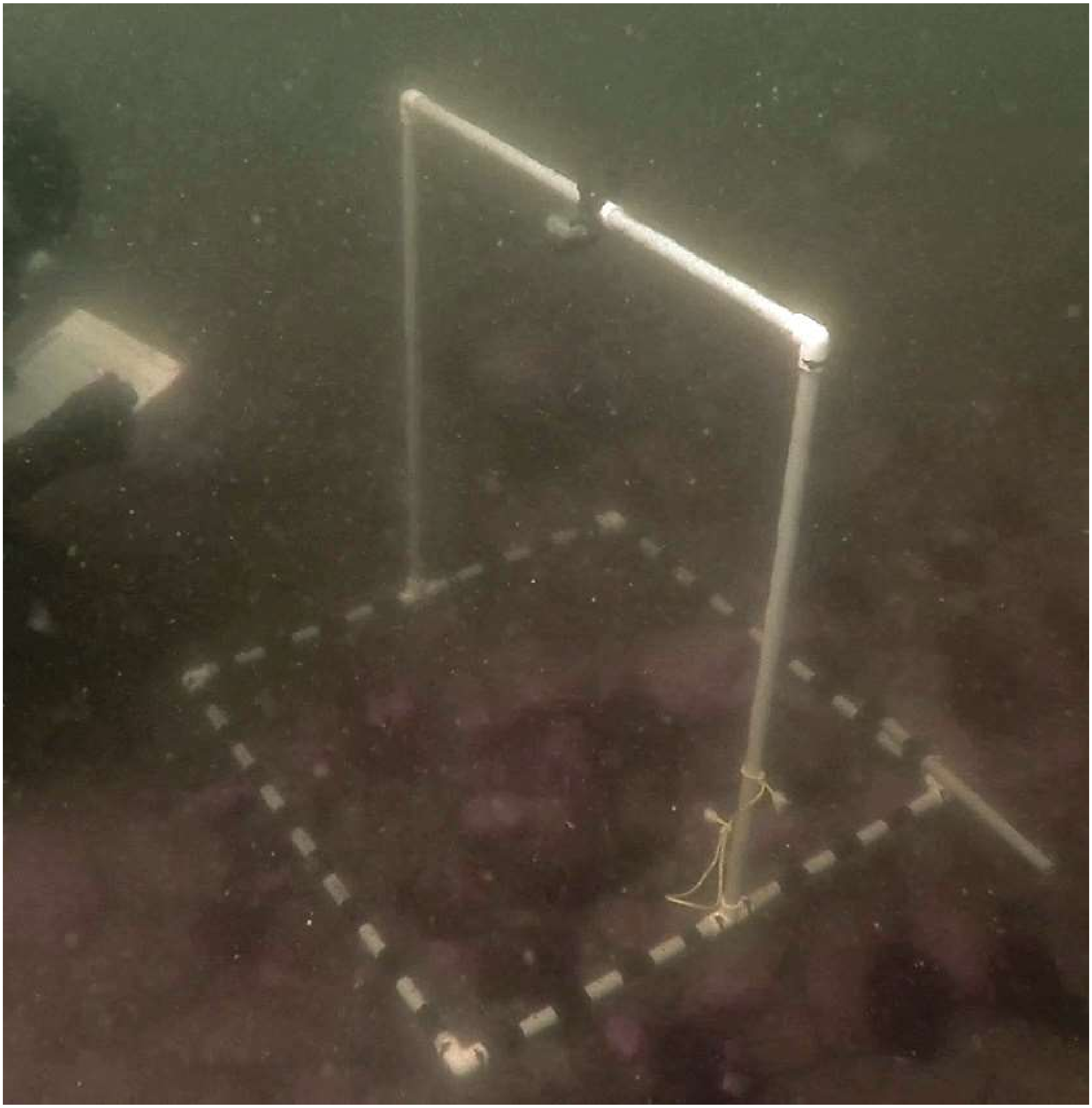

**Figure S2.** Foldable PVC frame for GoPro. The base of the frame is a quadrat with 80 cm long sides on which 5 cm reference markers have been applied using electrical tape.

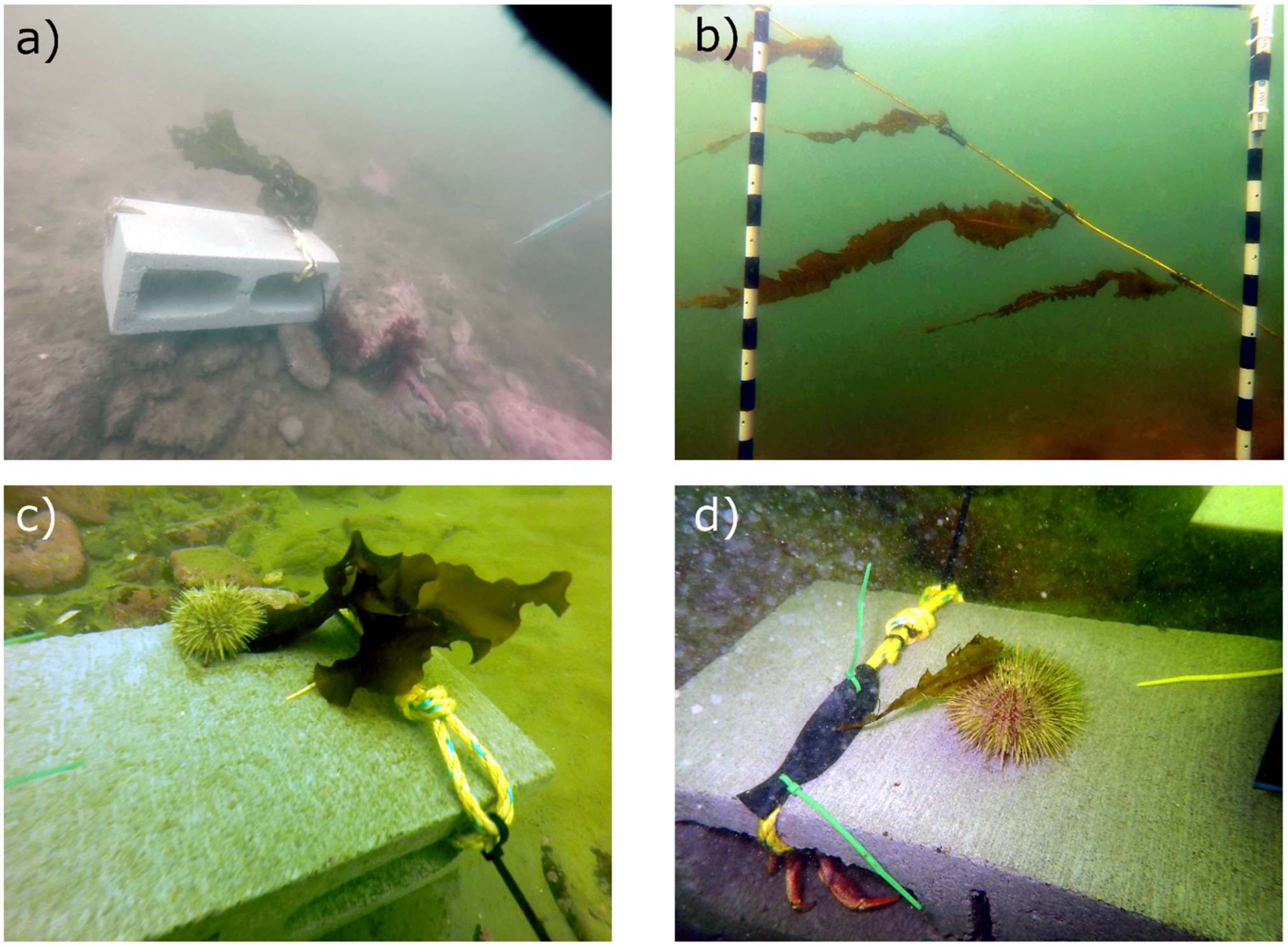


**Figure S3.** Representative pictures of the experiments**.** a) Each experimental unit consisted in a cinder block of 15×40×20 cm on which one individual of *Alaria esculenta* was attached by fixing the thallus on a rope using a rubber strip. b) Detail of the kelp frame showing kelp spaced on one line. c) and d) Example of urchins on the top of blocks in contact with kelp transplant, likely during a feeding interaction. In d) a crab can be seen hiding in one cavity of the block.

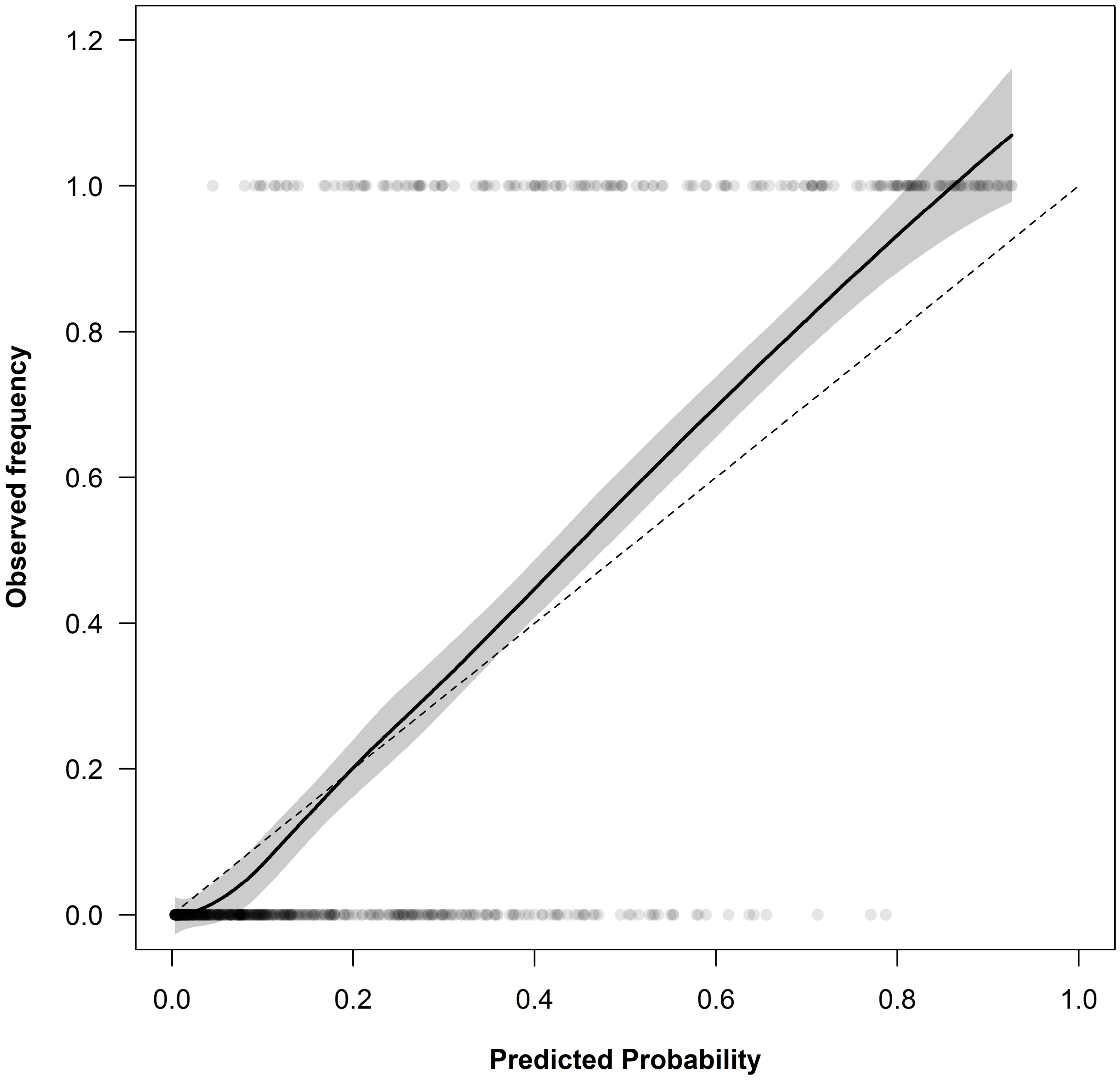


**Figure S4.** Calibration plot. Circles are drawn for each pair of observed response (urchin presence/absence in a cell in a given time) and its predicted probability. The dotted line represents the ideal case in which the frequency of observed responses with a given predicted probability is equal to the predicted probability (e.g. 20% of Y=1 responses have a predicted probability of 20%). The solid line represents the smoothing function describing the relationship between predicted and observed probabilities, while the gray shaded area represents its 0.95 confidence envelope. Where the smoother lies close to the diagonal, the model is well calibrated.

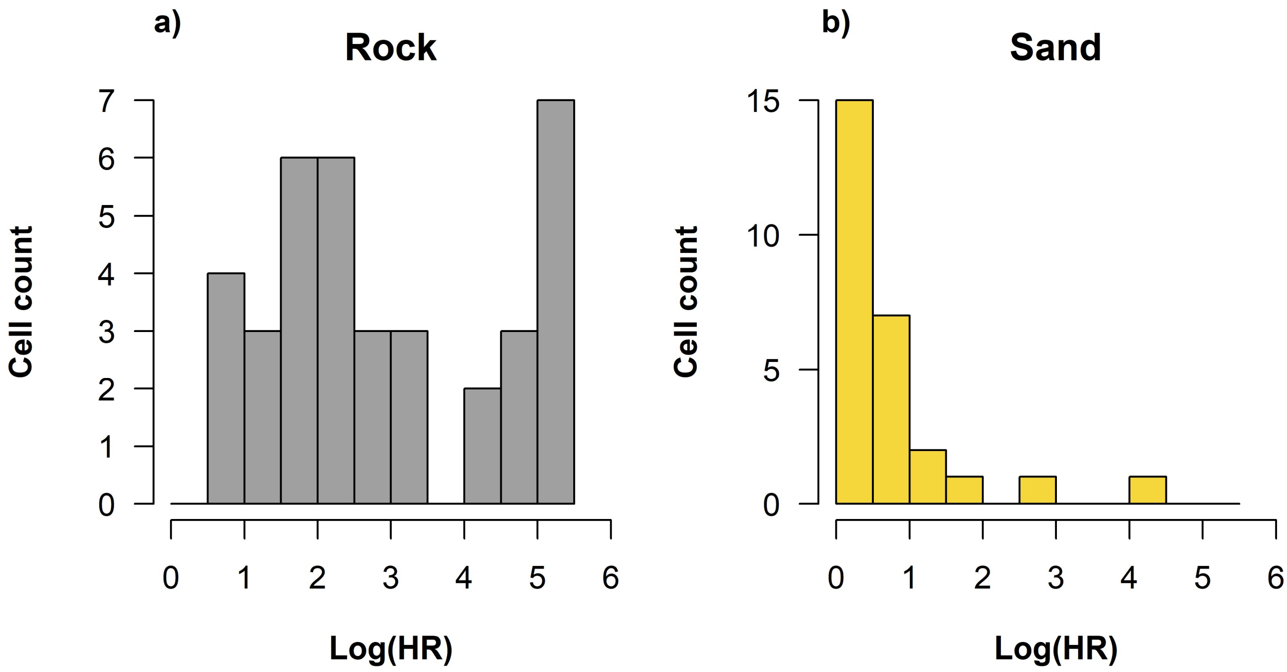


**Figure S5.** Distribution of grazing risk within rocky (a) and sandy (b) patch of the substrate. The histograms report the log hazard ration on the x-axis and the cell count on the y-axis. The grazing risk on the rocky patch is more uniformly distributed than on the sandy patch. The majority of cells in the sandy patch have the lowest level of risk.

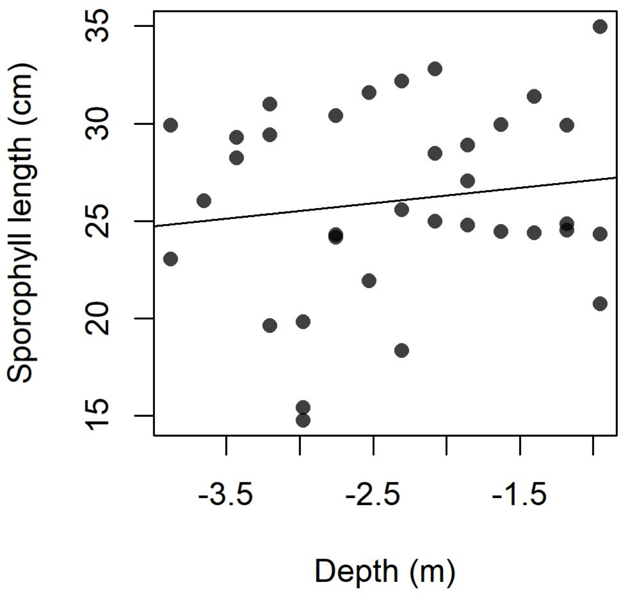


**Figure S6.** Length of sporophylls developed by transplant on the kelp frame. No significant effect of depth was found (P=0.4). Regression line equation is *length*=27.9 + 0.8 × *depth*.

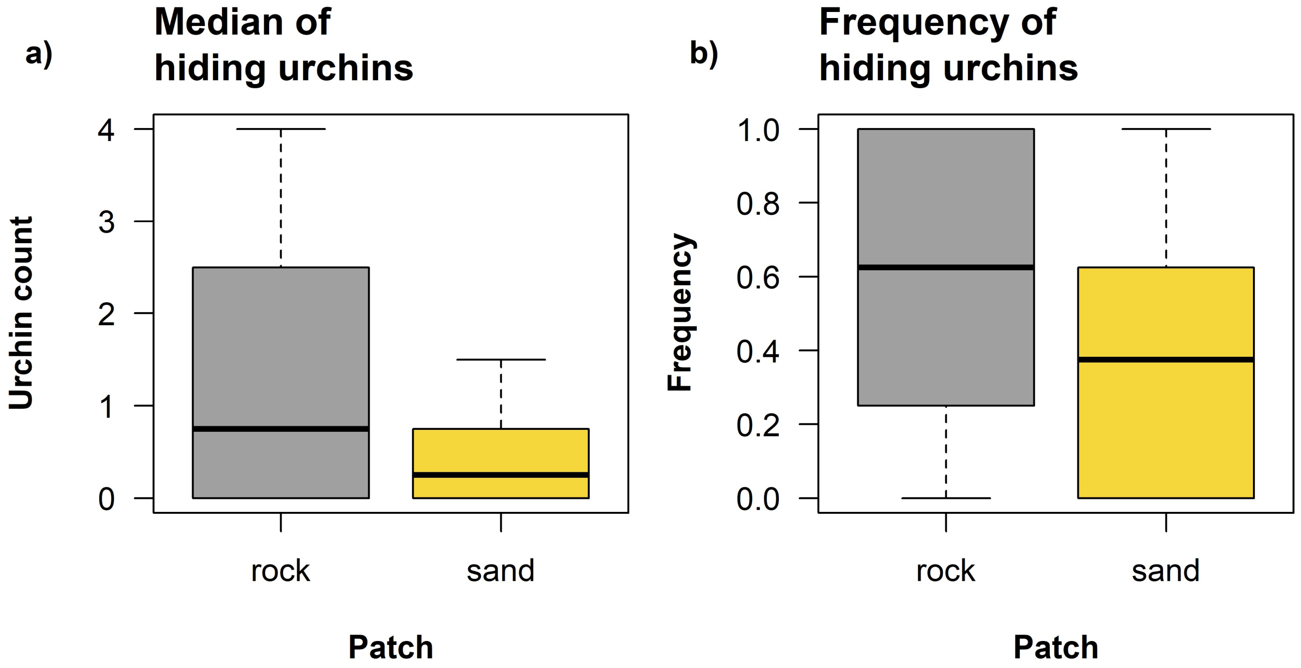


**Figure S7.** Differential use of the blocks' cavities by sea urchins depending on substrate type. The boxplots summarize a) the number of urchins found inside the blocks and b) the frequency with which urchins were found in the block, when blocks were on the rocky rather than the sandy patch.

### References


Barton, K. (2016). *MuMIn: Multi-Model Inference*. Retrieved from https://cran.r-project.org/package=MuMIn

Bates, D., Mächler, M., Bolker, B., & Walker, S. (2015). Fitting Linear Mixed-Effects Models Using lme4. *Journal of Statistical Software*, *67*(1), 1–48. doi: 10.18637/jss.v067.i01

Beijbom, O., Edmunds, P. J., Roelfsema, C., Smith, J., Kline, D. I., Neal, B. P., … Kriegman, D. (2015). Towards Automated Annotation of Benthic Survey Images: Variability of Human Experts and Operational Modes of Automation. *PLoS ONE*, *10*(7), e0130312. Retrieved from http://dx.doi.org/10.1371%2Fjournal.pone.0130312

Carney, L. T., Waaland, J. R., Klinger, T., & Ewing, K. (2005). Restoration of the bull kelp Nereocystis luetkeana in nearshore rocky habitats. *Marine Ecology Progress Series*, *302*, 49–61. doi: 10.3354/meps302049

Hijmans, R. J. (2016). *raster: Geographic Data Analysis and Modeling*. Retrieved from https://cran.r-project.org/package=raster

Kleinbaum, D. G., & Klein, M. (2005). *Survival analysis a self-learning text*. Retrieved from http://ariane.ulaval.ca/cgi-bin/recherche.cgi?qu=i9780387291505

Therneau, T. M. (2019). *survival: Survival Analysis*. Retrieved from https://cran.r-project.org/package=survival

Wentworth, C. K. (1922). A scale of grade and class terms for clastic sediments. *Journal of Geology*, *30*(5), 377–392.

Zuur, A. F., Ieno, E. N., Walker, N. J., Saveliev, A. A., & Smith, G. M. (2009). Mixed effects models and extensions in ecology with R. In M. Gail, K. Krickeberg, J. M. Samet, A. Tsiatis, & W. Wong (Eds.), *Statistics for Biology and Health*. doi: 10.1007/978-0-387-87458-6_1
